# Structure-Based Design, Synthesis and Biological Evaluation of Peptidomimetic Aldehydes as a Novel Series of Antiviral Drug Candidates Targeting the SARS-CoV-2 Main Protease

**DOI:** 10.1101/2020.03.25.996348

**Authors:** Wenhao Dai, Bing Zhang, Xia-Ming Jiang, Haixia Su, Jian Li, Yao Zhao, Xiong Xie, Zhenming Jin, Jingjing Peng, Fengjiang Liu, Chunpu Li, You Li, Fang Bai, Haofeng Wang, Xi Chen, Xiaobo Cen, Shulei Hu, Xiuna Yang, Jiang Wang, Xiang Liu, Gengfu Xiao, Hualiang Jiang, Zihe Rao, Lei-Ke Zhang, Yechun Xu, Haitao Yang, Hong Liu

## Abstract

SARS-CoV-2 is the etiological agent responsible for the COVID-19 outbreak in Wuhan. Specific antiviral drug are urgently needed to treat COVID-19 infections. The main protease (M^pro^) of SARS-CoV-2 is a key CoV enzyme that plays a pivotal role in mediating viral replication and transcription, which makes it an attractive drug target. In an effort to rapidly discover lead compounds targeting M^pro^, two compounds (**11a** and **11b**) were designed and synthesized, both of which exhibited excellent inhibitory activity with an IC50 value of 0.05 μM and 0.04 μM respectively. Significantly, both compounds exhibited potent anti-SARS-CoV-2 infection activity in a cell-based assay with an EC50 value of 0.42 μM and 0.33 μM, respectively. The X-ray crystal structures of SARS-CoV-2 M^pro^ in complex with **11a** and **11b** were determined at 1.5 Å resolution, respectively. The crystal structures showed that **11a** and **11b** are covalent inhibitors, the aldehyde groups of which are bound covalently to Cys145 of M^pro^. Both compounds showed good PK properties in *vivo*, and **11a** also exhibited low toxicity which is promising drug leads with clinical potential that merits further studies.

## Introduction

In late December 2019, a cluster of pneumonia cases caused by a novel coronavirus was reported in Wuhan, China^1,2,3^. Genomic sequencing showed that this pathogenic coronavirus is 96.2% identical to a bat coronavirus and shares 79.5% sequence identify to SARS-CoV^4,5,6^. This novel coronavirus was named as severe acute respiratory syndrome coronavirus 2 (SARS-CoV-2) by the International Committee on Taxonomy of Viruses, and the pneumonia was designated as COVID-19 by the World Health Organization (WHO) on February 11, 2020^7^. The epidemic spread rapidly to all provinces of China and to more than 159 countries and was announced as a global health emergency by WHO^8^. To make matters worse, no clinically effective vaccines or specific antiviral drugs are currently available for the prevention and treatment of COVID-19 infections. The combination of α-interferon and the anti-HIV drugs Lopinavir/Ritonavir (Kaletra®) is the current clinical treatment strategy, but the curative effect remains very limited and toxic side effects cannot be ignored. Remdesivir, a broad-spectrum antiviral drug developed by Gilead Sciences, Inc., is the clinical drug under development for the treatment of new coronavirus pneumonia, but more data are needed to prove its efficacy to treat COVID-19^9,10,11^. Specific anti-SARS-CoV-2 drugs with efficiency and safety are urgently needed.

A maximum likelihood tree based on the genomic sequence showed that the virus falls within the subgenus *Sarbecovirus* of the genus *Betacoronavirus*^6^. Coronaviruses are enveloped, positive-sense, single-stranded RNA viruses that lack the ability to correct errors occurring during RNA replication. This property results in high variability of CoVs and mutations that occur frequently and quickly under environmental and evolutionary stress. Therefore, RNA CoVs are highly prevalent and severe pathogens of viral diseases^12^. The genomic RNA of CoVs is approximately 30 k nt in length with a 5’-cap structure and 3’-poly-A tail, which has the largest viral RNA genome known to date and contains at least 6 open reading frames (ORFs). There is an a-1 frameshift between ORF1a and ORF1b which are the first ORF, about two-third of genome length, directly translating polyprotein (pp) 1a/1ab. These polyproteins will be processed by a 3C-like protease (3CL^pro^), also named as the main protease (M^pro^), and one or two papain-like proteases (PLPs) into 16 non-structural proteins (nsps). Subsequently, these nsps catalyze the synthesis of a nested set of subgenomic RNAs which are used as templates to directly translate main structural proteins including envelope (E), membrane (M), spike (S), and nucleocapsid (N) proteins. Therefore, these proteases, especially 3CL^pro^, play a vital role in the life cycle of coronavirus^13,14,15,16^.

3CL^pro^ (M^pro^) is a three-domain (domains I to III) cysteine protease and involves in most maturation cleavage events within the precursor polyprotein. Active 3CL^pro^ is a homodimer containing two protomers. The CoV 3CL^pro^ features a non-canonical Cys…His dyad located in the cleft between domains I and II^17,18,19^. Several common features are shared among the substrates of CoVs 3CL^pro^, and especially a Gln residue is almost absolutely required for the substrate in the P1 position. 3CL^pro^ is conserved within the group of CoVs. In addition, there is no human homologue of 3CL^pro^ which makes it an ideal antiviral target^20,21^.

### Design and synthesis of a series of peptidomimetic aldehydes as coronavirus 3CL protease inhibitors

The substrates of coronaviruses 3CL^pro^ (M^pro^) show some similarity, and most 3CL protease inhibitors are peptidomimetic covalent inhibitors derived from the natural substrates. The active sites are highly conserved among all CoV M^pro^ and are usually composed of four pockets (S1’, S1, S2 and S4) ^22,23^. The thiol of a cysteine residue in the S1’ pocket can anchor inhibitors by a covalent linkage, which is important for the inhibitors to maintain anti-viral activity. In our design of new inhibitors, the aldehyde was selected as a new warhead in P1’ to occupy the S1’pocket. As the (*S*)-γ-lactam ring has been proved to be suitable in the S1 pocket of 3C^pro^ and 3CL^pro^, this ring was expected to be a good choice in P1 of new inhibitors. Furthermore, the S2 pocket of coronavirus 3CL^pro^ is usually large enough to accommodate the bigger P2 fragment. To assess the possibility of π-π stacking interactions and hydrophobic interaction with the S2 pocket, the aryl and cyclohexyl group were placed in P2 (compounds **11a** and **11b**). Finally, the indole or other heterocyclice groups, which are privileged skeletons, were introduced into P3 in order to form new hydrogen bonds with S4 and improve drug-like properties.

Synthetic procedures: The synthetic route and chemical structures of the compounds (**11a** and **11b**) are shown in **Scheme 1**. The starting material **1** was obtained from commercial suppliers and used without further purification to synthesize the key intermediate **3** according to the literature^24^. The intermediates **6a** and **6b** were synthesized from **4** and acid **5a**, **5b**. After the *t*-butoxycarbonyl group was removed from **6a** and **6b**, the intermediates **7a** and **7b** were obtained. Coupling compounds **7a** and **7b** with the acid **8** yielded the esters **9a**, **9b**. The peptidomimetic aldehydes **11a** and **11b** were approached *via* a two-step route in which the ester derivatives **9** were first reduced with NaBH4 to generate the primary alcohols **10a** and **10b**, which were subsequently oxidized into aldehydes **11a** and **11b** with Dess-Martin Periodinane (DMP).

**Scheme 1.**
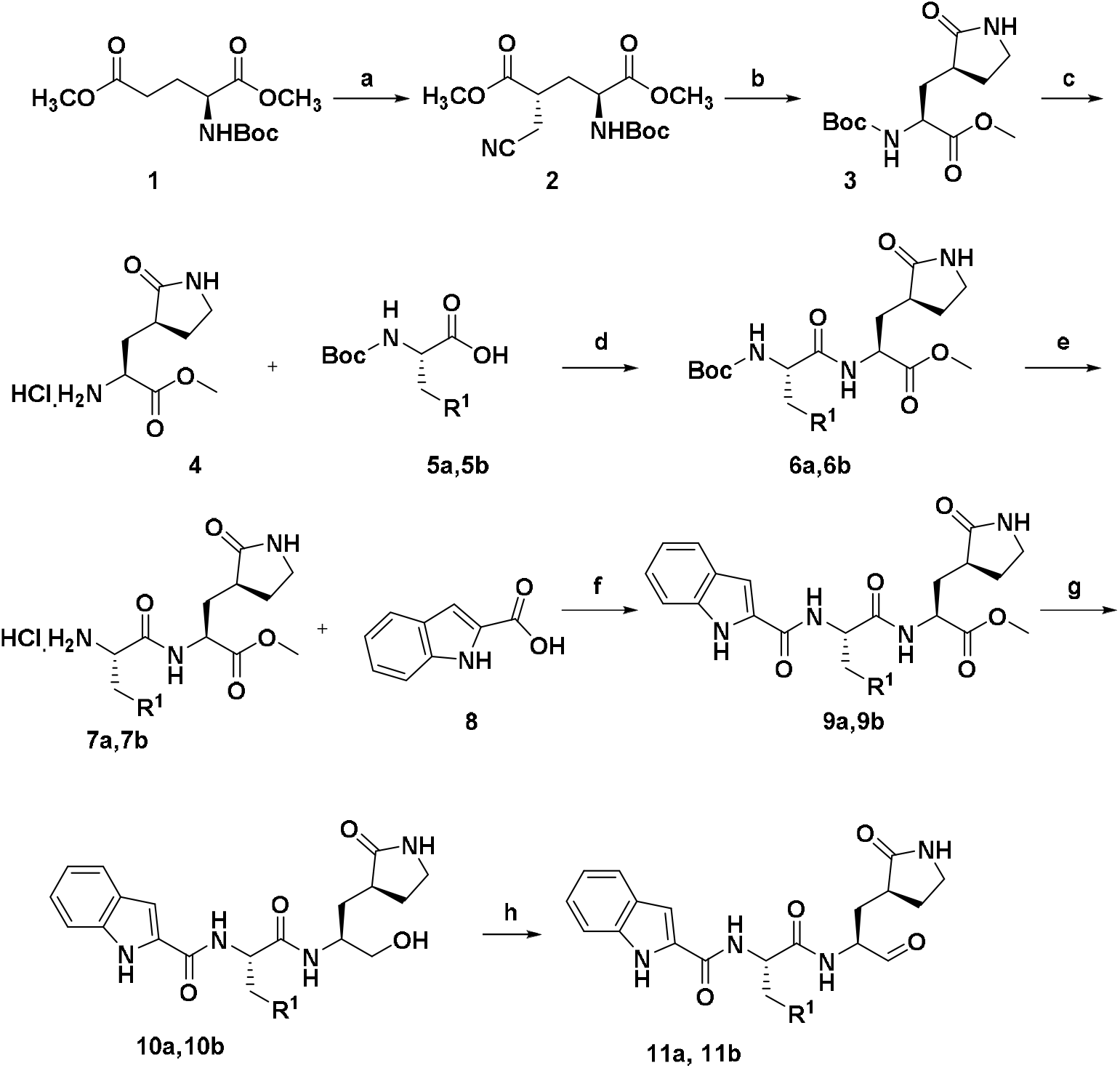
Reagents and Conditions: (a) LiHMDS, THF, −78°C; (b) NaBH_4_, CoCl_2_·6H_2_O, 0°C; (c) 4 M HCl, 12 h; (d) HATU, DIPEA, CH_2_Cl_2_, −20°C, 12 h; (e) 4 M HCl, 12 h; (f) HATU, DIPEA, CH_2_Cl_2_, −20°C, 12 h; (g) NaBH_4_, THF; (h) Dess-Martin Periodinane, CH_2_Cl_2_.

### Establishing a SARS-CoV-2 M^pro^ activity assay

Recombinant SARS-CoV-2 M^pro^ (3CL^pro^) was expressed and purified from *Escherichia coli (E. coli*)^18,25^. A fluorescently labeled substrate, MCA-AVLQ↓SGFR-Lys (Dnp)-Lys-NH_2_, derived from the *N*-terminal auto-cleavage sequence from the viral protease was designed and synthesized for the enzymatic assay.

Encouragingly, both compounds **11a** and **11b** exhibited high SARS-CoV-2 3CL^pro^ inhibition activity, which reached 100.4% for **11a** and 96.3% for **11b** at 1 μM, respectively. Further experiments were conducted by a fluorescence resonance energy transfer (FRET)-based cleavage assay to determine the IC_50_s. The results revealed excellent inhibitory potency with an IC_50_ value of 0.053 ± 0.005 μM and 0.040 ± 0.002 μM, respectively.

### The crystal structure of SARS-CoV-2 M^pro^ in complex with 11a

In order to elucidate the mechanism of inhibition of SARS-CoV-2 M^pro^ by **11a**, we determined the high-resolution crystal structure of this complex at 1.5-Å resolution (**Table S1**)^18,26,27^. The crystals of M^pro^-**11a** belongs to the space group *C2* and each asymmetric unit contains only one molecule (**Table S1**). By a crystallographic 2-fold symmetry axis, two molecules (designated protomer A and protomer B) associate into a homodimer (**Figure S2**). The structure of each protomer contains three domains and the substrate-binding site is located in the cleft between domain I and II (**Figure S2**). At the active site of SARS-CoV-2 M^pro^, Cys145 and His41 (Cys-His) form a catalytic dyad (**Figure S2**).

The electron density map clearly showed the compound **11a** in the substrate binding pocket of SARS-CoV-2 M^pro^ in an extended conformation (**Figure 2A and S3A**). To facilitate the explanation of the binding mode of **11a**, we will introduce it according to the chemical skeleton of this compound (P1’: aldehyde group; P1: (*S*)-γ-lactam ring; P2: cyclohexyl; P3: indole group). The electron density showed that the C of the aldehyde group of **11a** and the catalytic site Cys145 of SARS-CoV-2 M^pro^ form a standard 1.8 Å C–S covalent bond (**Figure 2B**), which suggests a Michael addition reaction. Furthermore, the oxygen atom of the aldehyde group also plays a crucial role for stabilizing the conformations of the inhibitor by forming hydrogen bonds with backbone of residues Cys145 and Gly143 in the S1’ site (**Figure 2B**). The (*S*)-γ-lactam ring of **11a** at P1 favorably inserts into the S1 site (**Figure 2B**). The oxygen of the (*S*)-γ-lactam group interacts with the side chain of His163 by hydrogen bond. The main chain of Phe140 and side chain of Glu166 also participate in stabilizing the (*S*)-γ-lactam ring by forming hydrogen bonds with the NH group. In addition, the amide bonds on the chain of **11a** are hydrogen-bonded with the main chains of His164 and Glu166, respectively (**Figure 2B**). The cyclohexyl moiety of 11a at P2 enters deep into the S2 site, stacking to the imidazole ring of His41 (Figure 2B). The cyclohexyl group is also surrounded by the side chains of Met49, Tyr54, Met165 and Asp187, producing extensive hydrophobic interactions (Figure 2B). The indole group of 11a at P3 is exposed to solvent (S4 site) and is stabilized by Glu166 through a hydrogen bond (Figure 2B). The side chains of residues Pro168 and Gln189 interact with the indole group of 11a through hydrophobic interactions. Interestingly, multiple water molecules (named W1-W6) play an important role in binding 11a (Figure 2B). W1 interacts with the amide bonds of 11a through a hydrogen bond, whereas W2-6 form a number of hydrogen bonds with the aldehyde group of 11a and the residues of Asn142, Gly143, Thr26, Thr25, His41 and Cys44, which contributes to stabilize 11a in the binding pocket (Figure 2B).

**Figure 1.**
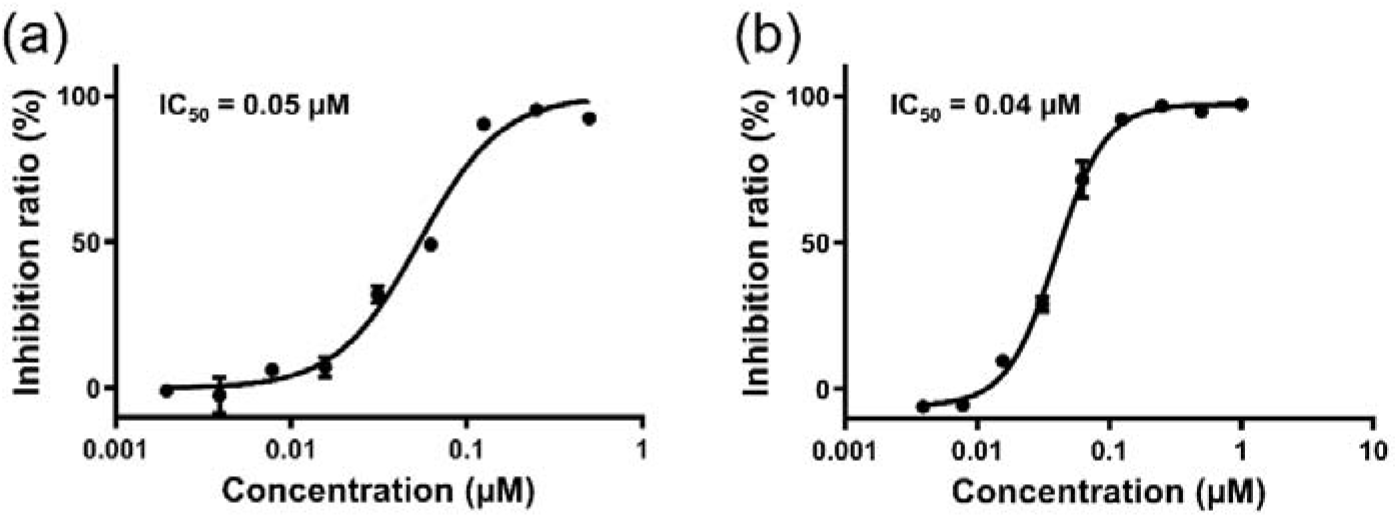
Inhibitory activity profiles of compounds **11a (a)** and **11b (b)** against SARS-CoV-2 M^pro^.

**Figure 2.**
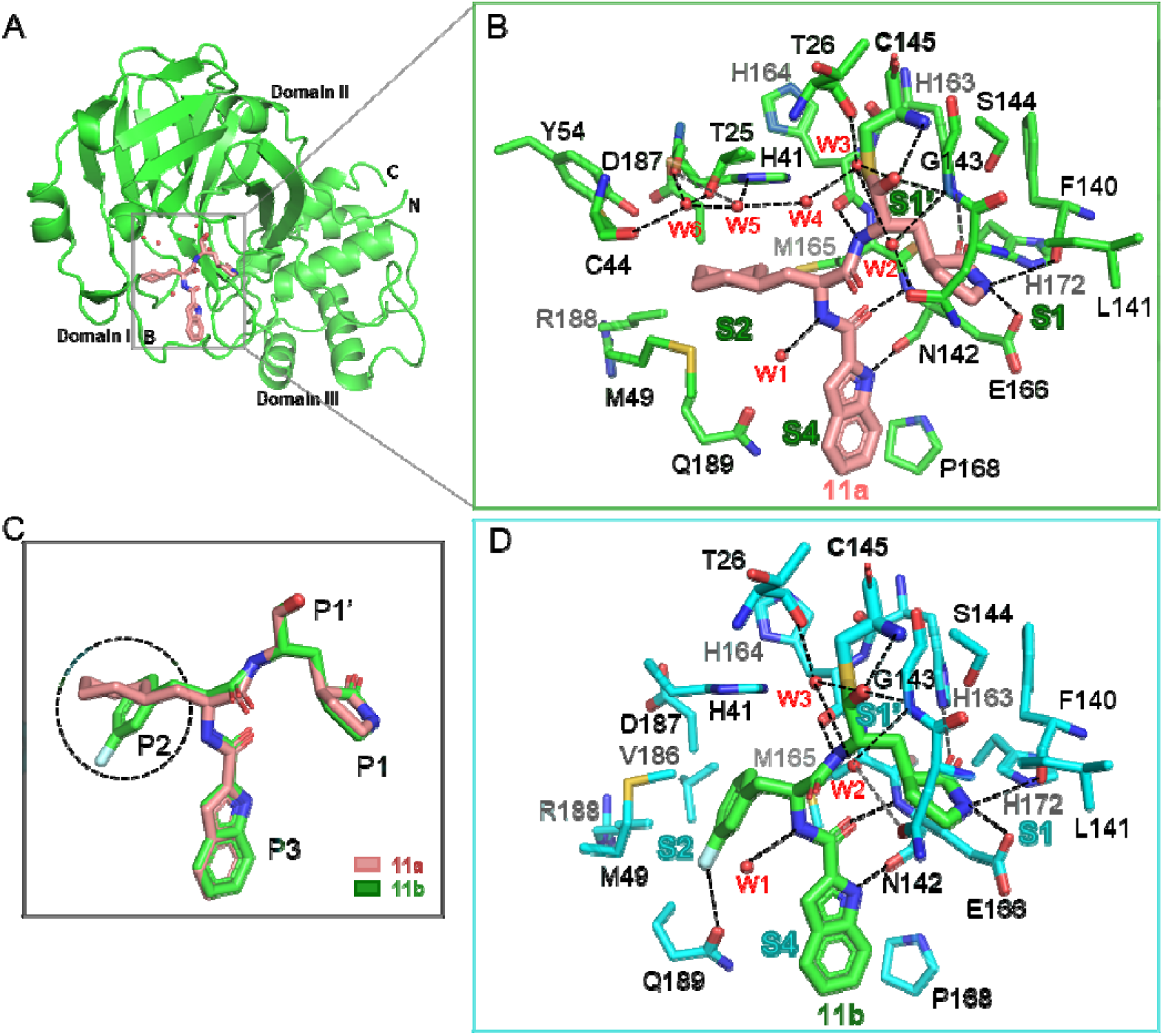
SARS-CoV-2 M^pro^ inhibitor binding pocket for 11a and 11b. A. Cartoon representation of the crystal structure of M^pro^ in complex with **11a**. The compound **11a** is shown as brown sticks in the substrate-binding pocket located between domain I and II of SARS-CoV-2 M^pro^. Water molecules involve in stabilizing the **11a** shown as spheres colored red. B. Close-up view of the **11a** binding site. The binding pocket is divided into four subsites (S1’, S1, S2 and S4). The residues involving in inhibitor binding are shown as green sticks. **11a** and water molecules are shown as brown sticks and red spheres, respectively. Hydrogen bonds are indicated as dashed lines. C. Comparison of the binding model of **11a** and **11b** in SARS-CoV-2 M^pro^. The major differences between **11a** and **11b** are marked with dashed circles. The compounds of **11a** and **11b** are shown as brown and green sticks, respectively. D. Close-up view of the **11b** binding site. Hydrogen bonds are indicated as dashed lines.

### The crystal structure of SARS-CoV-2 M^pro^ in complex with 11b

The crystal structure of SARS-CoV-2 M^pro^ in complex with **11b** is very similar to that of the **11a** complex and shows a similar inhibitor binding mode (**Figure 2C, 2D, S3B and S3C**). The difference in binding is probably due to the aryl group of **11b** at P2. Compared with the cyclohexyl group in **11a**, the aryl group undergoes a significant rotation (**Figure 2C**). The side chains of residues His41, Met49, Met165 and Val186 interact with this aryl group through hydrophobic interactions (**Figure 2D**). The side chain of Gln189 stabilizes the aryl group with an additional hydrogen bond (**Figure 2D**). In short, these two crystal structures reveal an identical inhibitory mechanism in that these two compounds occupy the substrate-binding pocket, mimicking the intermediates in the catalytic reaction, which blocks the enzyme activity of SARS-CoV-2 M^pro^.

### Antiviral activity assay

To further substantiate the enzyme inhibition results, we evaluated the ability of these compounds to inhibit SARS-CoV-2 *in vitro*. Vero E6 cells (ATCC-1586) were treated with a series of concentrations of the two compounds, and then were infected with a clinical isolate of SARS-CoV-2 (nCoV-2019BetaCoV/Wuhan/WIV04/2019) at a multiplicity of infection (MOI) of 0.05. At 24 hours post infection (h p.i.), viral copy numbers in the cell supernatant were quantified using quantitative real time PCR (RT-PCR). The cytotoxicity of these compounds in Vero E6 cells was also determined by using Cell Counting kit 8 (CCK8) assays. As shown in **Figure 3**, compounds **11a** and **11b** exhibited good anti-SARS-CoV-2-infection activity in cell culture with an EC50 values of 0.42 ± 0.08 μM and 0.33 ± 0.09 μM, respectively. Neither compound caused significant cytotoxicity, with half cytotoxic concentration (CC_50_) values of >100 μM, yielding a selectivity index (SI) of **11a** and **11b** of >238 and >303, respectively. Thus, **11a** and **11b** exhibit a very good antiviral effect on SARS-CoV-2.

**Figure 3.**
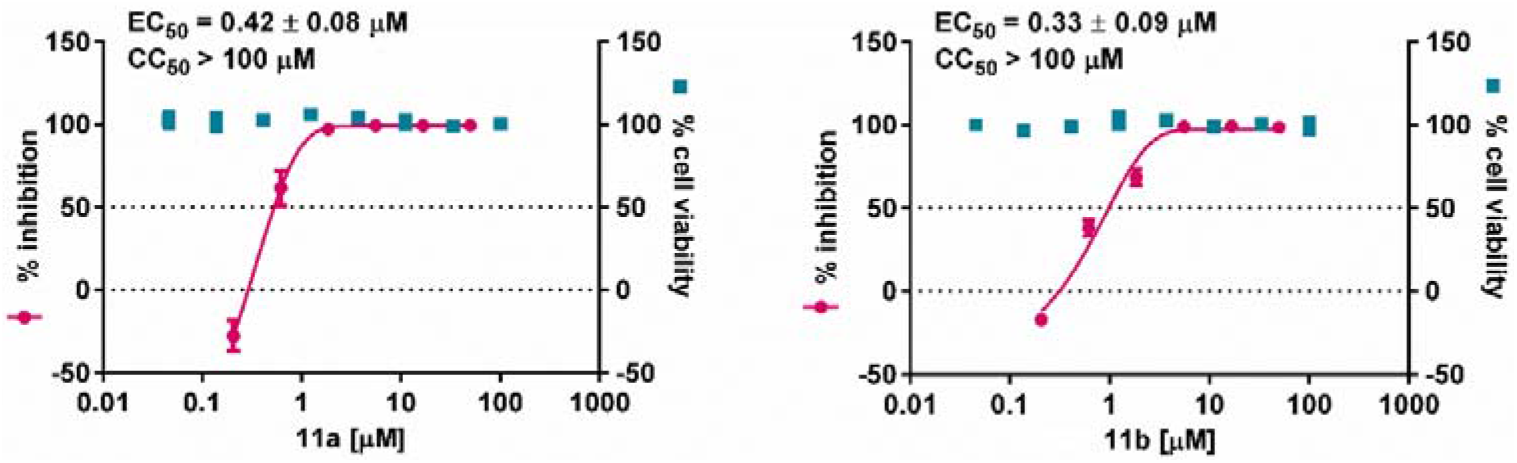
In vitro inhibition of viral 3CL protease inhibitors against SARS-CoV-2. Vero E6 cells were treated with a series concentration of indicated compounds **11a** and **11b** and infected with SARS-CoV-2 at an MOI of 0.05. At 24 hours post infection, cell supernatants were collected and the viral yield in the cell supernatant was quantified by qRT-PCR. The cytotoxicity of these compounds in Vero E6 cells was also determined by using CCK8 assays. The left and right Y-axis of the graphs represent mean % inhibition of virus yield and mean % cytotoxicity of the drugs, respectively.

### Preliminary pharmacokinetic (PK) evaluation of 11a and 11b

To explore the further druggability of the compounds **11a** and **11b**, both of two compounds were evaluated for its pharmacokinetic properties. As shown in **Table S2**, compound **11a** given intraperitoneally (5mg/kg) and intravenous (5mg/kg) displayed a long half-life (T_1/2_) of 4.27 h and 4.41h, a high maximal concentration (C_max_=2394 ng/mL), and a good bioavailability of 87.8%. Metabolic stability of **13a** in mice was also good (CL = 17.4 mL/min/mg). When administered intraperitoneal (20mg/kg), subcutaneous (5mg/kg) and intravenous (5mg/kg), compound **11b** also showed good PK properties. Considering the danger of COVID-19, we selected the intravenous drip administration to further study. Compared with **11a** administrated *via* intravenous, the half-life (1.65h) of **11b** is shorter and the clearance rate is faster (CL = 20.6 mL/min/mg). Compound **11a** was selected for further investigation with intravenous drip dosing on rats and dogs. The results showed (**Table S3)** that **11a** exhibited long T1/2 (rat, 7.6 h and dog, 5.5h), low clearance rate (rat, 4.01 mL/min/kg and dogs, 5.8 mL/min/kg) and high AUC value (rat, 41500 h*ng/mL and dog, 14900 h*ng/mL)). Those results indicating that compound **11a** has good PK properties to warrant further study.

### *In vivo* toxicity evaluation of 11a

The *in vivo* toxicity study (**Table S4)** of **11a** have been carried out on SD rats and Beagle dogs. The acute toxicity of **11a** was conducted on SD rats, and no SD rats died after receiving 40 mg/kg *via* intravenous drip administration. When the dosage was raised to 60 mg/kg, one of four SD rats was died. The dose range toxicity study of **11a** was conducted for seven days in the dosing level at 2, 6, 18 mg/kg on SD rats and at 10-40 mg/kg on Beagle dogs, once daily dosing (QD), by intravenous drip, all animals were clinically observed once a day at least and no obvious toxicity was observed in each group. The results show **11a** with low toxicity on rats and dogs.

## Discussion

New infectious agents have emerged to cause epidemics, such as SARS-CoV, MERS-CoV, and SARS-CoV-2. In order to identify antivirals to contain CoV infection, novel peptidomimetic aldehyde derivatives were designed, synthesized and evaluated biologically for their anti-SARS-CoV-2 main protease (M^pro^) activity and anti-SARS-CoV-2-infection activity in cell-based assays. Compounds **11a** and **11b** exhibited excellent anti-SARS-CoV-2 M^pro^ activity (IC50 = 0.053 ± 0.005 μM and IC_50_ = 0.040 ± 0.002 μM respectively) and good anti-SARS-CoV-2-infection activity in cell culture (EC_50_ = 0.42 ± 0.08 μM and EC_50_ = 0.33 ± 0.09 μM respectively). The crystal structures have shown that these drug leads can bind to the substrate-binding pocket of SARS-CoV-2 M^pro^, revealing the detailed covalent inhibition at the active site of the enzyme. Therefore, the class of peptidomimetic inhibitor carrying aldehydes has demonstrated potent inhibition both on the viral protease in the biochemical level and viral replication in the cell-based assays. Both compounds showed good PK properties in *vivo*, and **11a** also exhibited low toxicity which is promising compounds for heading to the clinical study.

## Supporting information

abstruct of paper

## Acknowledgment (Need supplement)

We thank Prof. James Halpert and LetPub (www.letpub.com) for its linguistic assistance during the preparation of this manuscript.

## Author contributions

H. Y. and H. L. conceived the project. Y. X., L. Z., H. Y., and H. L. designed the experiments; W.D. and J.L. designed and synthesized those compounds; X. X., J. P., C. L., S. H., J. W., performed the chemical experiments and collected the data. B. Z., Y. Z., Z.J., F. L., H. W., and X. Y. collected the diffraction data and solved the crystal structure; G. X., Z. R., Y X., H. J., H. Y., H. L.,Y. L., and Y. H. analyzed and discussed the data Y. X., L. Z., H. Y., and H. L., wrote the manuscript.

## Data and materials availability

All data are available in the main text or the supplementary materials. The PDB accession No. for the coordinates of SARS-CoV-2 M^pro^ in complex with **11a** is 6LZE, and the PDB accession No. for the coordinates of SARS-CoV-2 M^pro^ in complex with **11b** is 6M0K.

