## supporting information for "Structure-Based Design, Synthesis and Biological Evaluation of Peptidomimetic Aldehydes as a Novel Series of Antiviral Drug Candidates Targeting the SARS-CoV-2 Main Protease"

**Materials and Methods:**

**General procedures:** The materials and solvents were purchased from commercial sources and used without further purification. All products were characterized by their NMR and MS spectra. $^{1}$H and $^{13}$C NMR spectra were recorded on a 400 MHz, 500 MHz or 600 MHz instrument. Compounds were purified by chromatography with silica gel (300-400 mesh). Analytical thin layer chromatography (TLC) was HSGF 254 (0.15-0.2 mm thickness). High-resolution mass spectra (HRMS) were measured on Micromass Ultra Q-TOF spectrometer. All target compounds possessed a purity of ≥95% as determined by HPLC.

**Synthetic procedures:** The synthetic route and chemical structures of the compounds (**11a** and **11b**) were shown in Scheme S1. The starting material **1** was obtained from commercial suppliers and used without further purification to synthesize the key intermediate **3** by following the literature[24]. The intermediate **6a**, **6b** were synthesized by **4** and acid **5a**, **5b**. After the *t*-butoxycarbonyl group was removed from **6a**, **6b**, the intermediate **7a** and **7b** were obtained. Coupling compound **7a**, **7b** with the acids **8** yielded the corresponding **9a**, **9b**. The peptidomimetic aldehydes **11a** and **11b** were approached via a two-step route in which the ester derivatives **9a, 9b** were first reduced with $NaBH_4$ to generate the primary alcohols **10a** and **10b**, and then the alcohols **10a**, **10b** were subsequently oxidized into aldehydes **11a** and **11b** with Dess-Martin Periodinane (DMP).

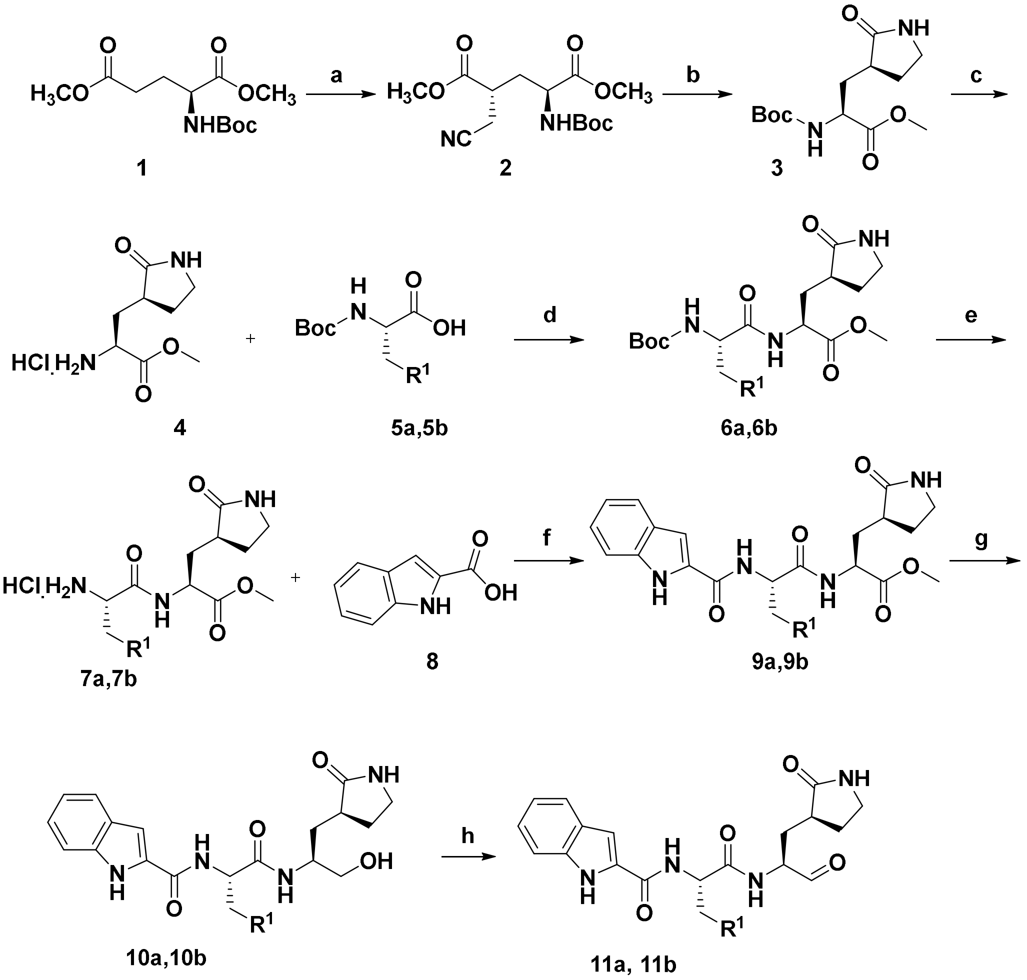


**Scheme S1.** Reagents and Conditions: (a) LiHMDS, THF, -78 °C; (b) $NaBH_4$, $CoCl_2{\cdot}6H_2O$, 0 °C; (c) 4 M HCl, 12 h; (d) HATU, DIPEA, $CH_2Cl_2$, -20 °C, 12 h; (e) 4 M HCl, 12 h ; (f) HATU, DIPEA, $CH_2Cl_2$, -20 °C, 12 h; (g) $NaBH_4$, THF; (h) Dess-Martin Periodinane, $CH_2Cl_2$.

**(2*S*,4*R*)-dimethyl 2-(*tert*-butoxycarbonylamino)-4-(cyanomethyl) pentanedioate (2)**

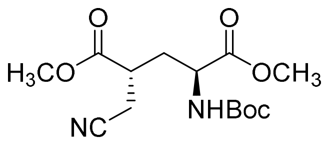

The solution of lithium bis(trimethylsilyl)amide (LHMDS) (94 mL, 1 M in THF) was added dropwise to a solution of *N*-Boc-*L*-glutamic acid dimethyl ester (12.0 g, 43.6 mmol) in THF (100 mL) at -78 °C, then the mixture was stirred at -78 °C for 1 h. Subsequently, bromoacetonitrile (3.24 mL, 46.6 mmol) was added dropwise to the mixture under the temperature of -78 °C, and the reaction was kept at -78 °C for additional 4 h. After the reactant was consumed, the reaction was quenched by $NH_4Cl$ (40 mL). The reaction mixture was warm up to room temperature and extracted with ethyl acetate (50 mL×3). The organic layers were concentrated and purified by flash column chromatography (petroleum ether/ethyl acetate = 4/1) to give product 2 (7.58 g, 55%) as colorless oil.

ESI-MS *m/z* 215.1 $(M - Boc + H)^+$.

$^1$H NMR (600 MHz, $CDCl_3$) δ 5.11 (d, *J* = 7.5 Hz, 1H), 4.38 (s, 1H), 3.77 (s, 3H), 3.75 (s, 3H), 2.92-2.82 (m, 1H), 2.81-2.71 (m, 2H), 2.24-2.08 (m, 2H), 1.44 (s, 9H).

**(*S*)-methyl 2-(*tert*-butoxycarbonylamino)-3-((*S*)-2-oxopyrrolidin-3-yl)propanoate (3)**

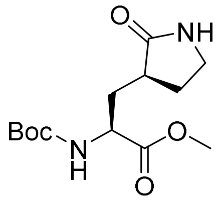


In a round-bottomed flask, the compound **2** (6.0 g, 19.09 mmol) was dissolved in anhydrous MeOH (100 mL) before $CoCl_2 \cdot 6H_2O$ (2.72 g, 11.45 mmol) was added at 0 °C. Subsequently, $NaBH_4$ (4.35 g, 114.78 mmol) was added protion-wise, and the reaction mixture was warm up to room temperature and stirred for 12 h. After the reactant was consumed, the reaction was quenched by $NH_4Cl$ (30 mL). MeOH in the mixture was evaporated and the residual mixture was extracted with ethyl acetate (50 mL×3). The organic layers were washed by saturated $NH_4Cl$ solution (100 mL×3) and brine (100 mL×3), then the organic phase was dried ($MgSO_4$) and concentrated. The residue was purified by flash column chromatography (petroleum ether/ethyl acetate = 2/1) to give the product 3 (2.18g, 40%) as white solid.

ESI-MS *m/z* 187.7 $(M - Boc + H)^+$.

$^1$H NMR (600 MHz, $CDCl_3$) δ 6.64 (s, 1H), 5.56 (s, 1H), 4.29 (d, *J* = 9.1 Hz, 1H), 3.71 (s, 3H), 3.37- 3.26 (m, 2H), 2.47-2.42 (m, 2H), 2.13-2.08 (m, 1H), 1.84-1.81 (m, 2H), 1.41 (s, 9H).

**methyl (*S*)-2-amino-3-((*S*)-2-oxopyrrolidin-3-yl)propanoate hydrochloride (4)**

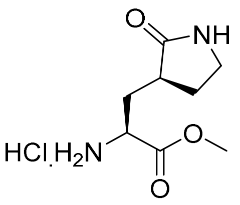


Compound **3** (1.0 g, 3.5 mmol) was dissolved in 10 mL DCM, then the HCl (9 mL, 4M in dioxane) was added. The reaction mixture was stirred at 20°C for 12 h, and the mixture was concentrated in vacuo to get a white solid **4**, which could be used for the following step without purification.

**methyl (*S*)-2-((*S*)-2-((*tert*-butoxycarbonyl)amino)-3-cyclohexylpropanamido)-3-((*S*)-2-oxopyrrolidin-3-yl)propanoate (6a)**

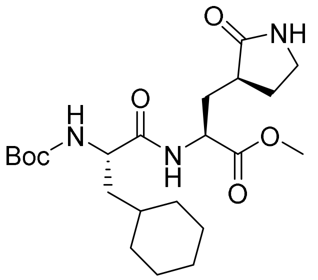


To the solution of Boc-*L*-Cyc-OH **5a** (0.95 g, 3.5 mmol) in DCM (40 mL) was added the HATU sequentially (1.9 g, 4.9 mmol) at -20 °C. The solution was kept at -20 °C for 20 mins, and then the crude product **4** (0.77 g 3.5 mmol) was added. After 30 min later, DIPEA (1.7 mL, 10.5 mmol) was added dropwise. And the reaction mixture was stirred at -20 °C for 12 h. The reaction mixture was washed by saturated $NH_4Cl$ solution (100 mL×3), saturated $NaHCO_3$ solution (100 mL×3) and brine (100 mL×3). The organic phase layer was dried over $Na_2SO_4$ and concentrated in vacuo. The resulting residue was purified by flash column chromatography (DCM: $CH_3OH$, 40: 1 v/v) to afford the pure product **6a** (1.23 g, 80%) as a light solid.

MS (ESI+) *m/z* 440.2 $(M+H)^+$.

$^1$H NMR (600 MHz, DMSO-$d_6$) δ 8.29 (d, *J* = 8.1 Hz, 1H), 7.60 (s, 1H), 6.83 (d, *J* = 8.1 Hz, 1H), 4.39-4.28 (m, 1H), 3.97-3.93 (m, 1H), 3.60 (s, 3H), 3.13 (t, *J* = 9.0 Hz, 1H), 3.06-3.04 (m, 1H),

2.36-2.24 (m, 1H), 2.11-2.02 (m, 2H), 1.70-1.55 (m, 7H), 1.47-1.38 (m, 1H), 1.35 (s, 9H), 1.32-1.19 (m, 2H), 1.17-1.04 (m, 3H), 0.85-0.81 (m, 2H).

**methyl (*S*)-2-((*S*)-2-amino-3-cyclohexylpropanamido)-3-((*S*)-2-oxopyrrolidin-3-yl) propanoate hydrochloride (7a)**

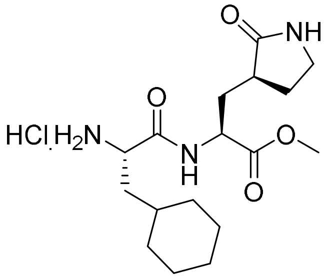


To a solution of **6a** (1.05 g, 2.4 mmol) in dry DCM was added the HCl (6 mL, 4M in dioxane) and the reaction mixture was stirred at ambient temperature for 12 h. Solvent was removed in vacuo and the crude product **7a** was directly used in next step without further purification.

**methyl (*S*)-2-((*S*)-3-cyclohexyl-2-(1*H*-indole-2-carboxamido)propanamido)-3-((*S*)-2-oxopyrrolidin-3-yl)propanoate (9a)**

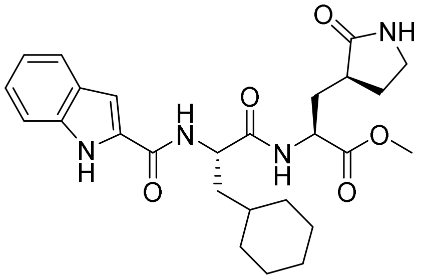


To a solution of the indole-2-carboxylic acid **8** (0.78 g, 2.4 mmol) in DCM was added the HATU (1.09 g, 2.88 mmol) sequentially at -20 °C. The solution was kept at -20 °C for 20 mins, and then the crude product **7a** (0.90 g 2.4 mmol) was added. DIPEA (1.17 mL, 7.2 mmol) was added dropwise after 30 mins later. Then, the reaction mixture was stirred at -20 °C for 12 h, followed by washing with saturated $NH_4Cl$ solution (100 mL×3), saturated $NaHCO_3$ solution (100 mL×3) and brine (100 mL×3). The organic phase was dried over $Na_2SO_4$ and concentrated, and the residue was purified by column chromatography ($CH_2Cl_2$: $CH_3OH$, 30: 1 v/v) to afford the pure product **9a** (0.98 g, 85%) as a light solid.

MS (ESI+) *m/z* 483.1 $(M+H)^+$.

$^1$H NMR (400 MHz, DMSO-$d_6$) δ 11.57 (s, 1H), 8.57 (d, *J* = 7.9 Hz, 1H), 8.42 (d, *J* = 8.0 Hz, 1H), 7.64 (s, 1H), 7.62 (d, *J* = 8.0 Hz, 1H), 7.42 (d, *J* = 8.2 Hz, 1H), 7.26 (d, *J* = 1.5 Hz, 1H), 7.18 (t, *J* = 7.5 Hz, 1H), 7.03 (t, *J* = 7.5 Hz, 1H), 4.59-4.57 (m, 1H), 4.40-4.31 (m, 1H), 3.62 (s, 3H), 3.16-3.08 (m, 2H), 2.37-2.35 (m, 1H), 2.14-2.04 (m, 2H), 1.76-1.72 (m, 2H), 1.70-1.55 (m, 8H), 1.45-1.35 (m, 1H), 1.20-1.12 (m, 2H), 0.97-0.88 (m, 2H).

**_N_-((_S_)-3-cyclohexyl-1-(((_S_)-1-hydroxy-3-((_S_)-2-oxopyrrolidin-3-yl)propan-2-yl)amin- o)-1-oxopropan-2-yl)-1_H_-indole-2-carboxamide (10a)**

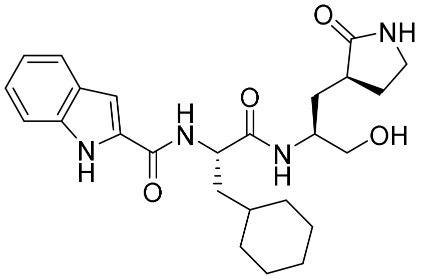


To a solution of **9a** (0.96 g, 2.0 mmol) in dry THF was added the $NaBH_4$ (0.45 g, 12 mmol) protion-wise at 0 °C and the $CH_3OH$ was added dropwise, then the reaction mixture was stirred at room temperature for 3 h. The completion of the reaction was confirmed by TLC then the reaction was quenched by $NH_4Cl$ solution (20 mL). The reaction mixture was extracted with ethyl acetate (50 mL×3) and the organic layers were washed with saturated $NH_4Cl$ solution (50 mL×3) and brine (50 mL×3). The organic phase was dried over $Na_2SO_4$ and concentrated, and the residue was purified by column chromatography (DCM: $CH_3OH$, 20: 1 v/v) to afford the pure product **10a** (0.82 g, 90%) as a light solid.

MS (ESI+) *m/z* 455.3 $(M+H)^+$.

$^1$H NMR (400 MHz, Methanol-$d_4$) δ 11.58 (s, 1H), 8.39 (d, J = 8.0 Hz, 1H), 7.78 (d, J = 9.0 Hz, 1H), 7.62 (d, J = 8.0 Hz, 1H), 7.52 (s, 1H), 7.42 (d, J = 8.2 Hz, 1H), 7.26 (d, J = 1.2 Hz, 1H), 7.17 (t, J = 7.5 Hz, 1H), 7.03 (t, J = 7.5 Hz, 1H), 4.68 (t, J = 5.6 Hz, 1H), 4.53-4.51 (m, 1H), 3.83-3.76 (m, 1H), 3.38-3.33 (m, 1H), 3.28-3.21 (m, 1H), 3.11 (t, J = 8.7 Hz, 1H), 3.05-3.01 (m, 1H), 2.30-

2.34 (m, 1H), 2.17-2.09 (m, 1H), 1.85-1.78 (m, 1H), 1.76-1.72 (m, 2H), 1.69-1.61 (m, 3H), 1.61-1.53 (m, 3H), 1.41-1.34 (m, 2H), 1.26 -1.12 (m, 2H), 1.04-0.78 (m, 3H).

***N*-((*S*)-3-cyclohexyl-1-oxo-1-(((*S*)-1-oxo-3-((*S*)-2-oxopyrrolidin-3-yl)propan-2-yl)amino) propan-2-yl)-1*H*-indole-2-carboxamide (11a).**

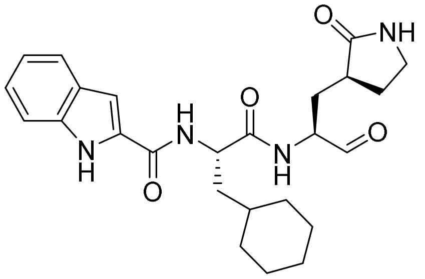


To a solution of the **10a** (0.45 g, 1.0 mmol) in DCM, the DMP (0.55 g, 1.2 mmol) was added slowly and the reaction mixture was stirred at room temperature. When the reactant was consumed, the reaction was filtered and washed with saturated $Na_2S_2O_3$ solution (50 mL×3), saturated $NaHCO_3$ solution (50 mL×3) and brine (50 mL×3). The organic phase was dried over $Na_2SO_4$ and concentrated, then the residue was purified by flash column chromatography (DCM: $CH_3OH$, 20: 1 v/v) to afford the pure product **11a** (0.34 g, 76%) as a white solid.

HRMS (*m/z*): $[M-H]^+$ calculated for $C_{25}H_{31}N_4O_4$, 451.2351; found, 451.2345.

HPLC purity： 99.88%. **11a** was determined by Agilent-1100 HPLC with binary pump, photodiode array detector (DAD), using Agilent Extend-C18 column (150 x 4.6 mm, 5 μm). **11a** was analyzed using MeOH/$H_2O$ = 65:35 (v/v) (30 min, 1 mL/min) and calculated the peak areas at 230 nm.

$^1$H NMR (500 MHz, DMSO) δ 11.58 (s, 1H), 9.44 (s, 1H), 8.59 (d, *J* = 7.6 Hz, 1H), 8.49 (d, *J* = 7.9 Hz, 1H), 7.68-7.57 (m, 2H), 7.44 (d, *J* = 8.2 Hz, 1H), 7.29 (d, *J* = 1.2 Hz, 1H), 7.19 (t, *J* = 7.5 Hz, 1H), 7.05 (t, *J* = 7.5 Hz, 1H), 4.64-4.59 (m, 1H), 4.25-4.20 (m, 1H), 3.20-3.06 (m, 2H), 2.37-2.30 (m, 1H), 2.18-2.13 (m, 1H), 1.96-1.92 (m, 1H), 1.81-1.58 (m, 9H), 1.42 (d, *J* = 2.9 Hz, 1H), 1.24-1.10 (m, 3H), 1.02-0.88 (m, 2H).

$^{13}$C NMR (126 MHz, DMSO) δ 201.30, 178.76, 173.52, 161.54, 136.93, 131.76, 127.51, 123.85, 122.01, 120.18, 112.73, 103.97, 56.85, 51.18, 39.89, 39.56, 37.79, 34.21, 33.65, 32.31, 29.78, 27.76, 26.56, 26.27, 26.11

***N*-((S)-3-(3-fluorophenyl)-1-oxo-1-(((S)-1-oxo-3-((S)-2-oxopyrrolidin-3-yl)propan-2-yl)amino)propan-2-yl)-1H-indole-2-carboxamide. (11b).**

The synthesis procedure of **11b** is similar as **11a**.

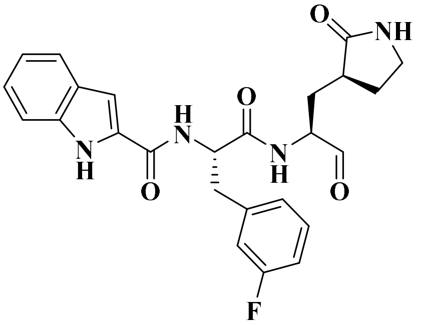


HRMS (ESI) m/z: calculated for $C_{25}H_{26}FN_4O_4^+$ $(M + H)^+$: 465.1933, found: 465.1936.

HPLC purity：99.20%. **11b** was determined by Agilent-1100 HPLC with binary pump, photodiode array detector (DAD), using Agilent Extend-C18 column (150 x 4.6 mm, 5 μm). **11b** was analyzed using MeOH/$H_2O$ = 65:35 (v/v) (20 min, 1 mL/min) and calculated the peak areas at 254 nm.

$^{1}$H NMR (500 MHz, Acetone) δ 10.92 (s, 1H), 9.44 (s, 1H), 8.57 (d, *J* = 6.8 Hz, 1H), 8.09 (d, *J* = 8.4 Hz, 1H), 7.60 (d, *J* = 8.0 Hz, 1H), 7.50 (dd, *J* = 8.3, 0.6 Hz, 1H), 7.28 (td, *J* = 7.9, 6.2 Hz, 1H), 7.22-7.11 (m, 5H), 7.07-7.03 (m, 1H), 6.93 (td, *J* = 8.3, 1.9 Hz, 1H), 5.08 (td, *J* = 8.4, 5.6 Hz, 1H), 4.38-4.34 (m, 1H), 3.36 (dd, *J* = 13.8, 5.6 Hz, 1H), 3.28-3.17 (m, 3H), 2.49-2.40 (m, 1H), 2.31-2.25 (m, 1H), 2.02-1.96 (m, 1H), 1.85-1.79 (m, 1H), 1.78-1.71 (m, 1H).

$^{13}$C NMR (126 MHz, Acetone) δ 200.80, 180.25, 172.63, 163.60 (d, *J* = 243.6 Hz), 162.40, 141.55 (d, *J* = 7.5 Hz), 137.87, 131.92, 130.89 (d, *J* = 8.3 Hz), 128.61, 126.37 (d, *J* = 2.7 Hz), 124.77,

122.63, 120.94, 117.03 (d, $J$ = 21.3 Hz), 114.13 (d, $J$ = 21.1 Hz) 113.17, 104.24, 58.57, 55.42, 40.89, 38.88, 38.37, 30.51, 29.10.

$^{19}$F NMR (376 MHz, Acetone) δ -113.99- -114.92 (m).

**Figure S1. Spectral data for compounds 11a and 11b.**

**$^{1}$H and $^{13}$C NMR, HRMS and HPLC spectra of 11a**

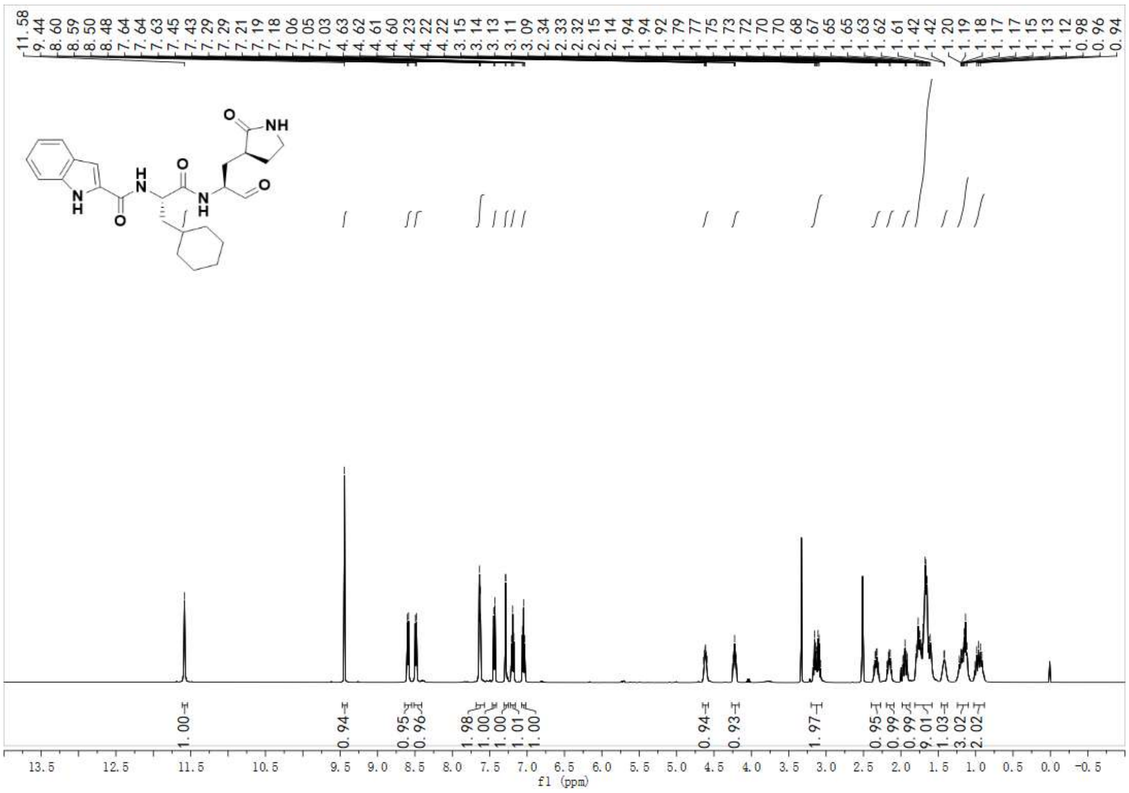

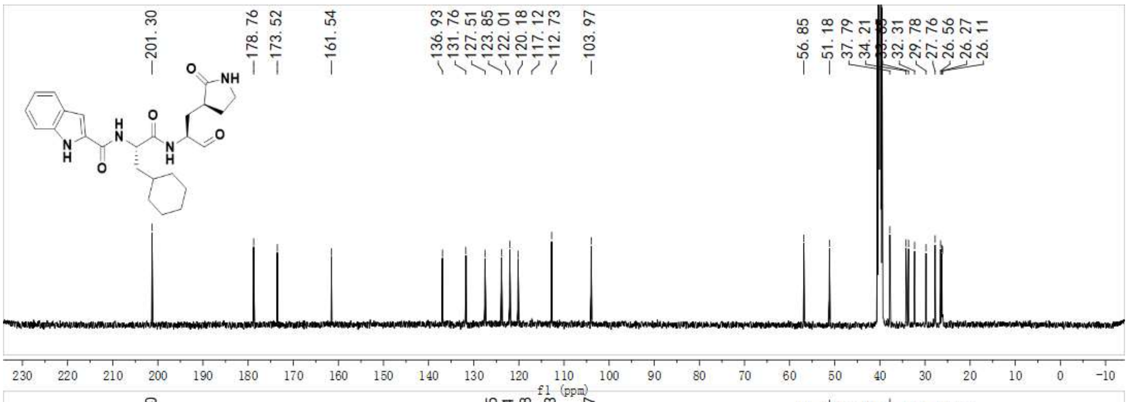


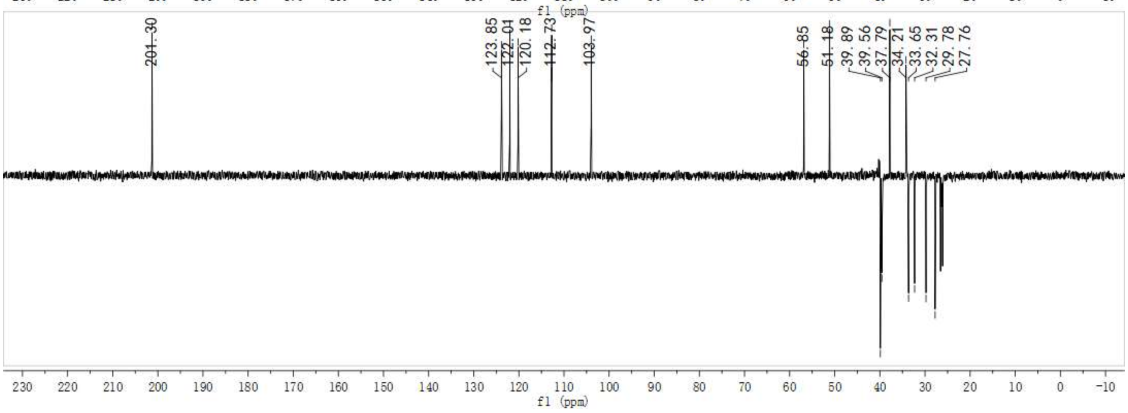


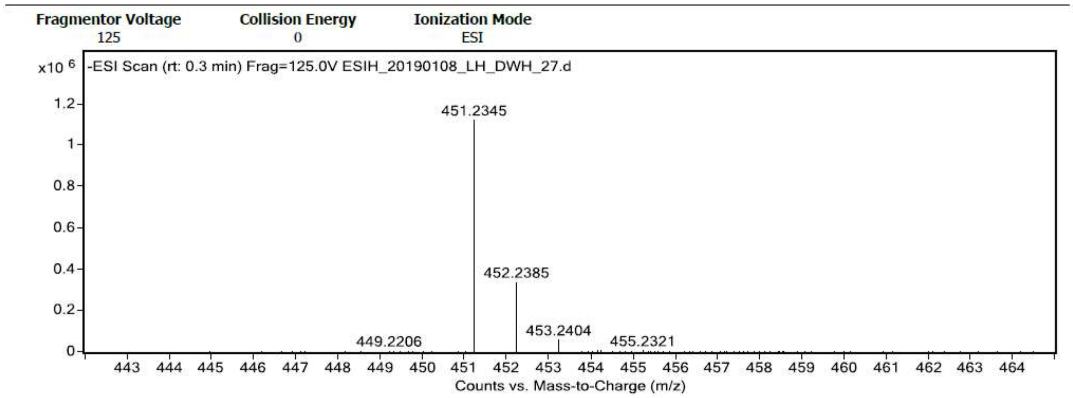


**Formula Calculator Results**

| m/z | Calc m/z | Diff (mDa) | Diff (ppm) | Ion Formula | Ion |
|---|---|---|---|---|---|
| 451.2345 | 451.2351 | 0.55 | 1.22 | C25 H31 N4 O4 | (M-H)- |

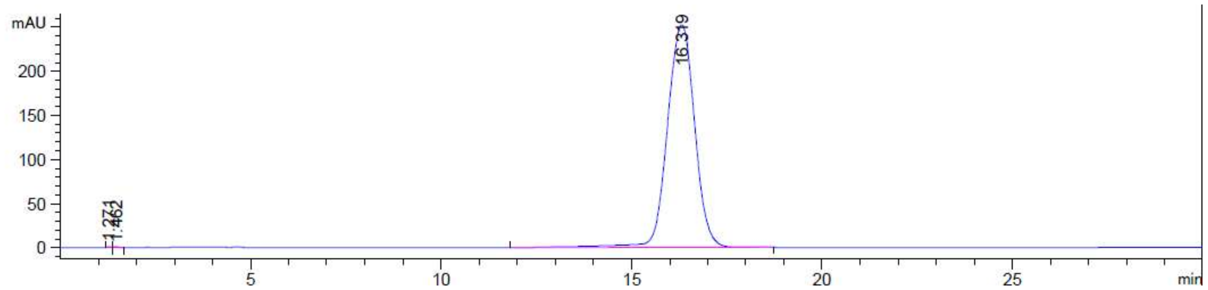

Signal 4: DAD1 D, Sig=230,4 Ref=360,100

| Peak # | RetTime [min] | Type | Width [min] | Area [mAU*s] | Height [mAU] | Area % |
|---|---|---|---|---|---|---|
| 1 | 1.271 | BV | 0.0634 | 6.19408 | 1.46259 | 0.0488 |
| 2 | 1.462 | VB | 0.1093 | 8.42439 | 1.23994 | 0.0664 |
| 3 | 16.319 | BB | 0.7941 | 1.26750e4 | 252.29979 | 99.8848 |
| Totals : | | | | 1.26896e4 | 255.00232 | |

**$^1$H, $^{13}$C and $^{19}$F NMR, HRMS and HPLC spectra of 11b.**

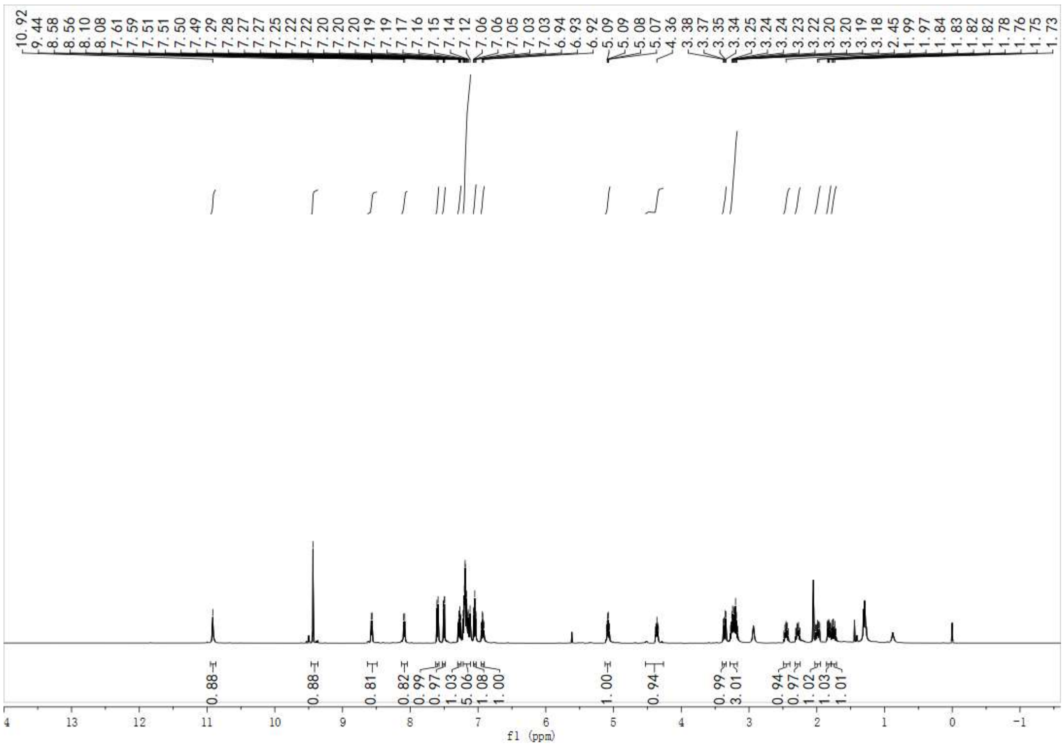


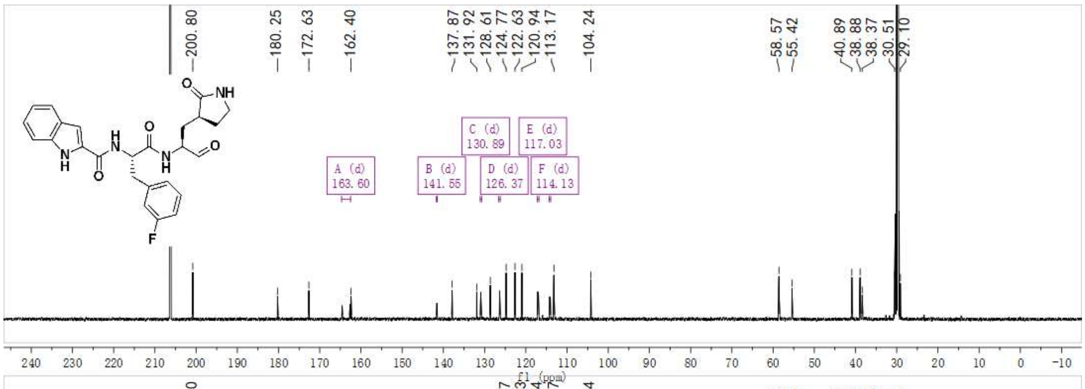


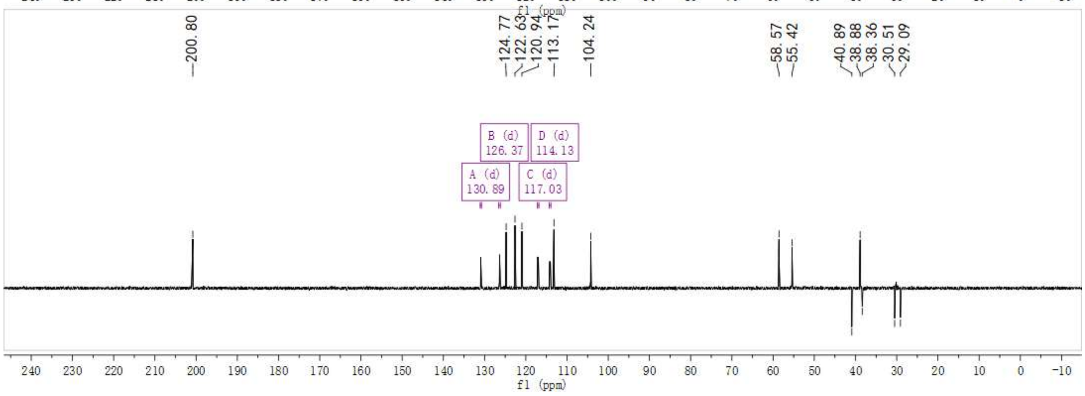

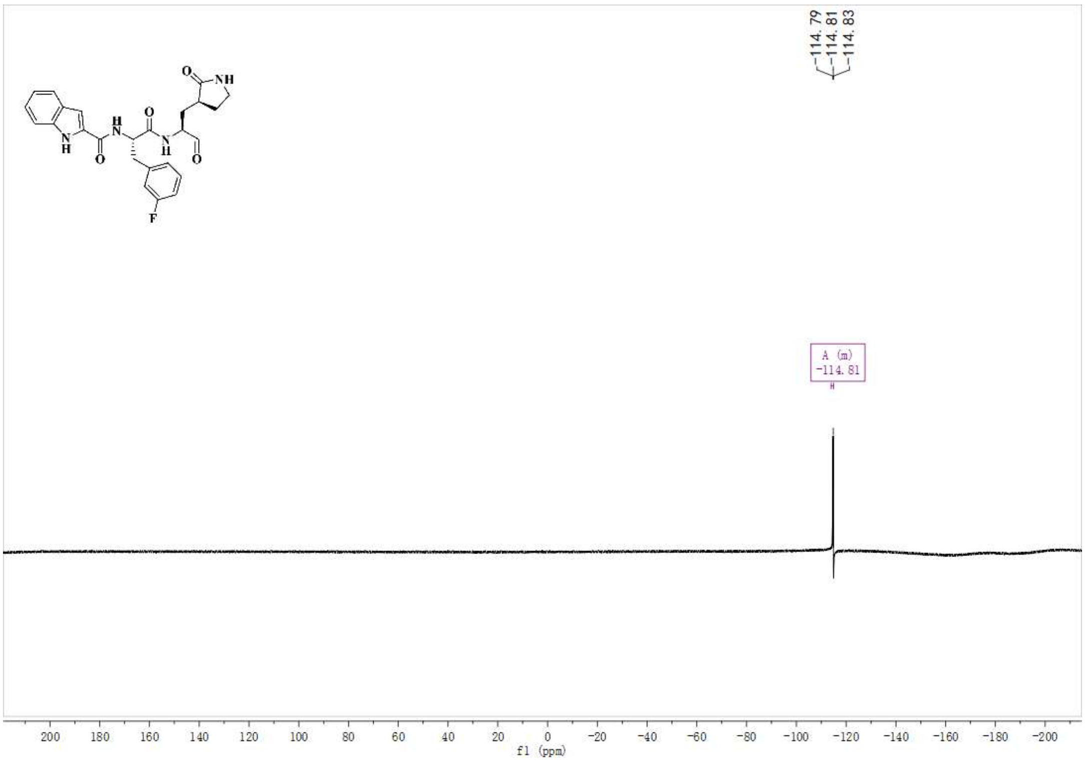


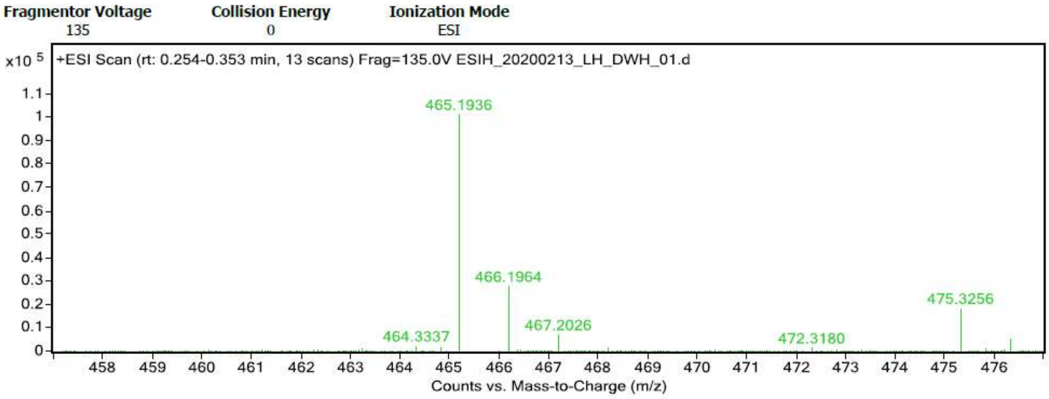


**Formula Calculator Results**

| m/z | Calc m/z | Diff (mDa) | Diff (ppm) | Ion Formula | Ion |
|---|---|---|---|---|---|
| 465.1936 | 465.1933 | -0.39 | -0.83 | C25 H26 F N4 O4 | (M+H)+ |

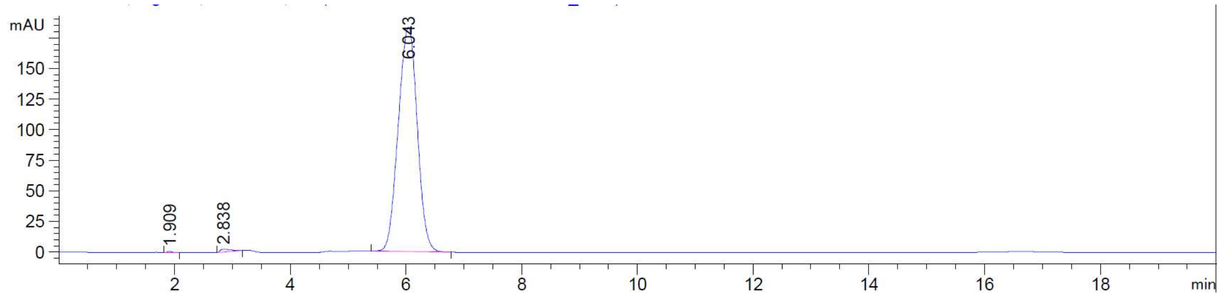

Signal 1: DAD1 A, Sig=254,4 Ref=360,100

| Peak # | RetTime [min] | Type | Width [min] | Area [mAU*s] | Height [mAU] | Area % |
|---|---|---|---|---|---|---|
| 1 | 1.909 | BB | 0.0766 | 5.42105 | 1.07946 | 0.1289 |
| 2 | 2.838 | BB | 0.1781 | 28.37407 | 2.36262 | 0.6745 |
| 3 | 6.043 | BB | 0.3637 | 4172.72705 | 183.22050 | 99.1966 |
| Totals : | | | | 4206.52217 | 186.66258 | |

**Cloning, expression and purification of** SARS-CoV-2 M$^{pro}$

The full-length gene encoding SARS-CoV-2 M$^{pro}$ was optimized and synthesized for *Escherichia coli* (*E. coil*) expression (GENEWIZ). The method of cloning and producing authentic SARS-CoV-2 M$^{pro}$ was followed by the protocol published for SARS-CoV M$^{pro}$ previously[18].

**Enzymatic activity and inhibition assays**

The enzyme activity and inhibition assays have been described previously.[22,25] The recombinant SARS-CoV-2 M$^{pro}$ (30 nM at a final concentration) was mixed with serial dilutions of each compound in 80 μL assay buffer (50 mM Tris–HCl, pH 7.3, 1 mM EDTA) and incubated for 10 min. The reaction was initiated by adding 40 μL fluorogenic substrate with a final concentration of 20 μM. After that, the fluorescence signal at 320 nm (excitation)/405 nm (emission) was immediately measured every 30 s for 10 min with a Bio-Tek Synergy4 plate reader. The *V*max of reactions added with compounds at various concentrations compared to the reaction added with DMSO were calculated and used to generate $IC_{50}$ curves. For each compound, $IC_{50}$ values against SARS-CoV-2 M$^{pro}$ were measured at 9 concentrations and three independent experiments were performed. All experimental data was analyzed using GraphPad Prism software.

**Crystallization**

SARS-CoV-2 M$^{pro}$ was incubated with 10 mM **11a** or **11b** for 30 min and the complex (5 mg/ml) was crystallized by hanging drop vapor diffusion method at 20 °C. The best crystals were grown with well buffer containing 2% polyethylene glycol (PEG) 6000, 3% DMSO, 1 mM DTT, 0.1 M MES (pH 6.0). The cryo-protectant solution contained 30% PEG 400, 0.1 M MES (pH 6.0).

**Data collection and structure determination**

All data were collected on beamline BL19U1 at Shanghai Synchrotron Radiation Facility (SSRF) at 100 K and at a wavelength of 0.9785 Å using an Pilatus3 6M image plate detector. Data integration and scaling were performed using the program XDS[18]. The structures were determined by molecular replacement (MR) with the SARS-CoV $M^{pro}$ (PDB ID: 2H2Z) as a search model using the paogram PHASER[26]. The output model from MR was subsequently subjected to iteative cycles of manual model adjustment with Coot[27] and refinement was finished with Phenix[18]. The inhibitor **11a** and **11b** was built according to the omit map. Data collection and structure refinement statistics are summarized in Table S1.

Coordinates and structure factors for SARS-CoV-2 $M^{pro}$ in complex with the inhibitor **11a** and **11b** have been deposited in Protein Data Bank with accession number 6LZE and 6M0K, respectively.

**Table S1. Data collection and refinement statistics**

| | **Mpro-11a** | **Mpro-11b** |
|---|---|---|
| **PDB code** | 6LZE | 6M0K |
| **Data collection** | | |
| Space group | *C*2 | *C*2 |
| Cell dimensions | | |
| *a, b, c* (Å) | 97.70, 80.94, 51.74 | 98.15, 81.70, 51.67 |
| *α, β, γ* (°) | 90, 114.27, 90 | 90, 114.69, 90 |
| Wavelength (Å) | 0.97852 | 0.97852 |
| Resolution (Å) | 50.00-1.51 (1.54-1.51) [a] | 50.00-1.50 (1.54-1.50) [a] |
| Completeness (%) | 98.0 (91.6) | 98.8 (90.3) |
| $R_{merge}$ (%) | 4.2 (55.5) | 3.0 (77.1) |
| Redundancy | 3.3 (2.7) | 3.4 (2.8) |
| $I/\sigma(I)$ | 13.99 (1.80) | 18.69 (1.30) |
| **Refinement** | | |

| | | |
|---|---|---|
| Resolution (Å) | 47.16-1.50 | 43.45-1.50 |
| No. of reflections | 57,378 | 58,412 |
| $R_{work}/R_{free}$ (%) | 17.8/20.1 | 18.34 / 19.66 |
| No. of atoms | | |
| Protein | 2340 | 2,347 |
| Ligand | 49 | 50 |
| Water | 209 | 163 |
| $B$ factor (Å$^2$) | | |
| Protein | 28.75 | 31.92 |
| Ligand | 37.60 | 52.56 |
| Water | 37.95 | 40.62 |
| R.m.s deviations | | |
| Bond lengths (Å) | 0.014 | 0.017 |
| Bond angles (°) | 1.280 | 1.440 |
| Ramachandran plot (%) | | |
| Favored | 98.0 | 98.0 |
| Allowed | 2.0 | 2.0 |
| Outliers | 0.0 | 0.0 |

[a] Values in parentheses are for highest-resolution shell.

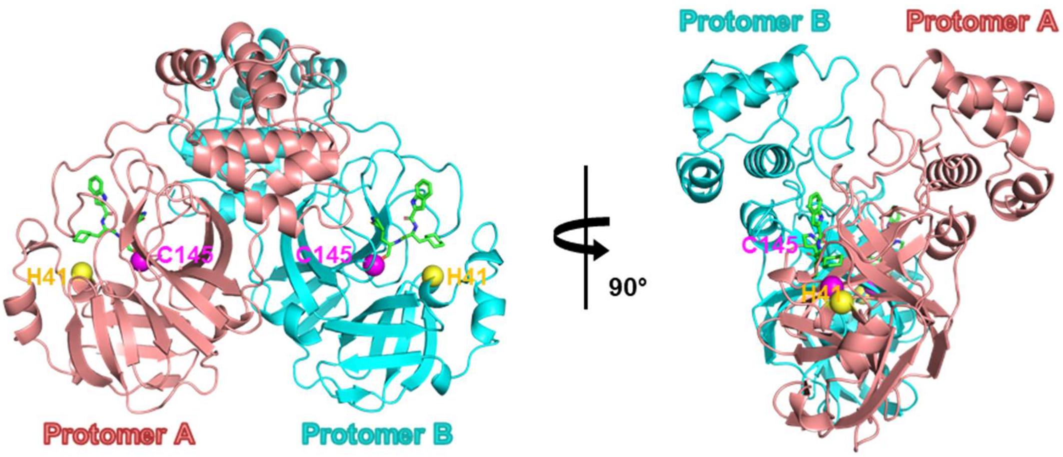


**Figure S2. Overall structure of SARS-CoV-2 M$^{pro}$**

Cartoon representation of SARS-CoV-2 $M^{pro}$ bound **11a** crystal structure in two different views. One protomer (protomer A) of the dimer is shown in brown, the other one (protomer B) in cyan. The residues of the catalytic site are indicated as yellow and magenta spheres, for His41 and Cys145, respectively. The compound **11a** is shown as green sticks.

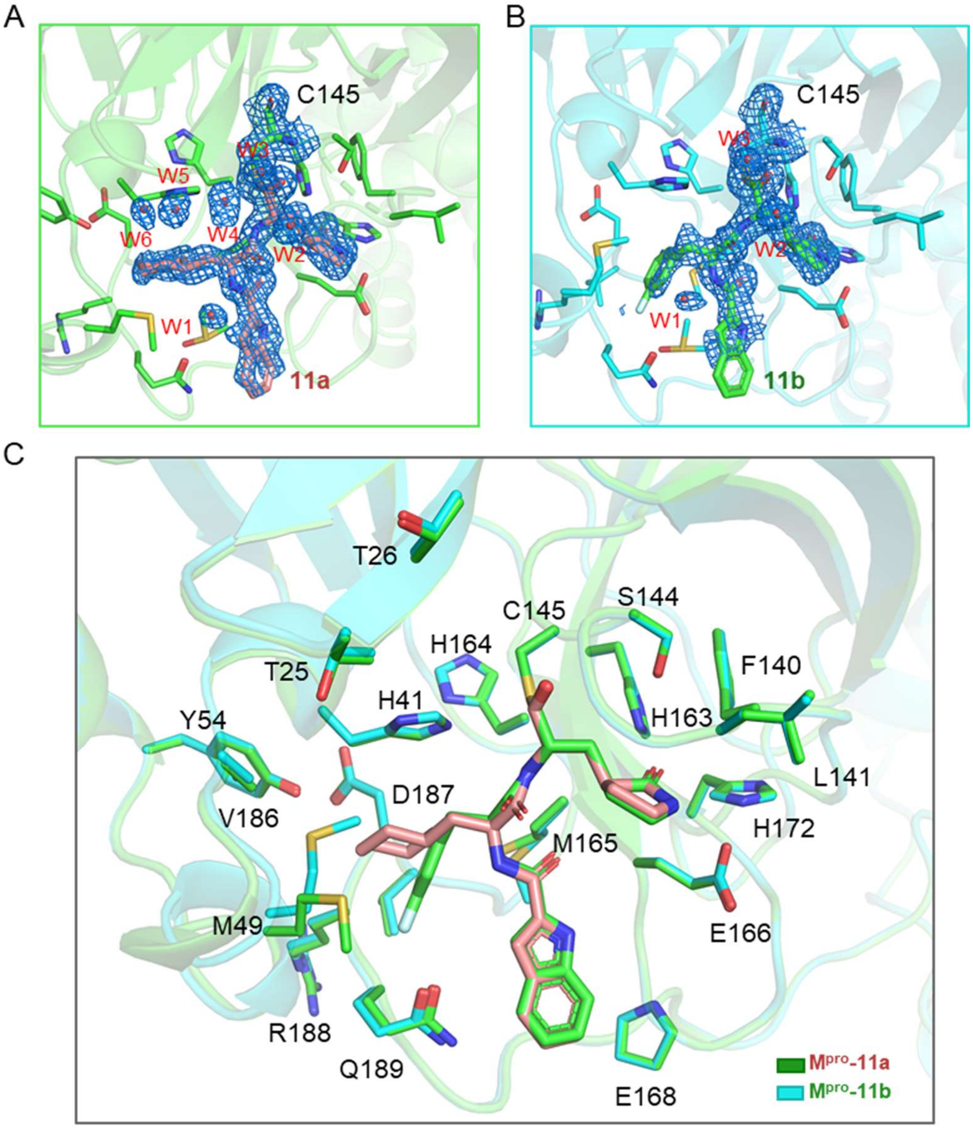


**Figure S3. Representative electron density of inhibitors and waters in the complex structures and Comparison of the 11a and 11b binding sites**

A, B. The $2F_o$-$F_c$ electron densities of **11a**, **11b**, the residues Cys145 and water molecules are shown as blue mesh and contoured at 1σ.

C. Comparison of 11a and 11b binding sites. Residues in 11a-bound and 11b-bound structures are colored green and cyan, respectively. **11a** and **11b** are shown as sticks colored brown and green, respectively.

**Antiviral assay for 11a and 11b**

Vero E6 cell line was obtained from American Type Culture Collection (ATCC) and maintained in Dulbecco's Modified Eagle Medium (DMEM; Gibco Invitrogen) supplemented with 10% fetal bovine serum (FBS; Gibco Invitrogen), 1% antibiotic/antimycotic (Gibco Invitrogen), at 37 °C in a humidified 5% $CO_2$ incubator. A clinical isolate SARS-CoV-2 was propagated in Vero E6 cells, and viral titer was determined as described previously[10]. All the infection experiments were performed at biosafety level-3 (BLS-3).

To assess the antiviral activity of compounds, pre-seeded Vero E6 cells ($5\times10^4$ cells/well) were treated with the different concentration of the indicated compounds for 1 hour, and then were infected with SARS-CoV-2 at a MOI of 0.05. Two hours later, the virus-drug mixture was removed and cells were further cultured with drug containing medium. At 24 h p.i., the cell supernatant was collected and viral RNA copy number was measured as described previously[10].

**Pharmacokinetics of 11a and 11b**

Male CD-1 mice (n = 3 per group) were treated with a solution of compounds **11a** and **11b** (DMSO/EtOH/PEG300/NaCl (5/5/40/50, v/v/v/v)) at doses of 20 mg/kg, 5 mg/kg and 5 mg/kg via intraperitoneal (ip), subcutaneous (sc) and intravenous (iv), respectively. Blood samples were collected at 0.05, 0.25, 0.75, 2, 4, 8, and 24 h after administration. Serum samples were obtained through common procedures and the concentrations of compound in the supernatant were analyzed by LC-MS/MS.

The solution of compound **11a** (HS15/NaCl (7.5/100, v/v)) was injected to SD rats and beagle dogs (n=2 per group) at doses of 10 mg/kg and 5 mg/kg via intravenous (iv). Blood samples were collected at 0.05, 0.5, 1, 2, 4, 6, 8, 10. 12 and 24 h after administration. Serum samples were obtained through common procedures and the concentrations of compound in the supernatant were analyzed by LC-MS/MS.

All procedures relating to animal handling, care, and treatment were performed according to the guidelines approved by the Institutional Animal Care and Use Committee of the contract research organizations performing the study.

**Table S2. Preliminary pharmacokinetic (PK) evaluation of compounds 11a and 11b in mice.[a]**

| Compd. | Admin | $T_{1/2}$ (h) | $T_{max}$ (h) | $C_{max}$ (ng/mL) | $AUC_{last}$ (h*ng/mL) | $AUC_{INF\_obs}$ (h*ng/mL) | CL (mL/min/kg) | $MRT_{INF\_obs}$ (h) | $Vss_{\_obs}$ (mL/kg) | F (%) |
|---|---|---|---|---|---|---|---|---|---|---|
| 11a | i.p(5mg/kg) | 4.27 ± 1.23 | 0.25 ± 0.00 | 2394 ± 288 | 4252 ± 367 | 4272 ± 366 | - | 1.85 ± 0.13 | | 87.8 |
| | i.v(5mg/kg) | 4.41 ± 0.30 | - | - | 4844 ± 757 | 4871 ± 766 | 17.4 ± 2.76 | 1.78 ± 0.19 | 1845 ± 211 | |
| 11b | i.p.(20mg/kg) | 5.21 ± 1.35 | 0.25 ± 0.00 | 12783 ± 421 | 13877 ± 2756 | 14090 ± 2677 | - | 2.23 ± 0.34 | | 85.9 |
| | s.c.(5mg/kg) | 2.22 ± 1.96 | 0.25 ± 0.00 | 3019 ± 665 | 3307 ± 939 | 3360 ± 973 | - | 1.41 ± 0.81 | | 81.8 |
| | i.v.(5mg/kg) | 1.65 ± 0.45 | - | - | 4041 ± 370 | 4076 ± 381 | 20.6 ± 2.0 | 0.63 ± 0.128 | 768 ± 96 | |

[a]The value of mice represented the average results from three independent experiments.

**Table S3. Preliminary pharmacokinetic (PK) evaluation of compound 11a in rat and dog.[a]**

| Compd. | Animal | Admin | $T_{1/2}$ (h) | $C_{3min}$ (ng/mL) | $AUC_{last}$ (h*ng/mL) | $AUC_{INF\_obs}$ (h*ng/mL) | CL (mL/min/kg) | $MRT_{INF\_obs}$ (h) | $Vss_{\_obs}$ (mL/kg) |
|---|---|---|---|---|---|---|---|---|---|
| 11a | Rat | i.v(10mg/kg) | 7.6± 4.8 | 81500 ± 700 | 41500 ± 200 | 41500 ± 100 | 4.01 ± 0.02 | 0.8 ± 0.00 | 203.5 ± 7.8 |
| | Dog | i.v(5mg/kg) | 5.5 ± 0.7 | 21900 ± 5700 | 14900 ± 4000 | 14900 ± 4000 | 5.80 ± 1.57 | 1.4 ± 0.1 | 489.5 ± 159.1 |

[a]The value of rat and dog represented the average results from two independent experiments.

### In *vivo* toxicity study of 11a

#### Acute toxicity study of 11a

SPF SD rats were half male and half female, which were weighted of 180~220 g for females and 210~250 g for males. Acute toxicity studies were performed in GLP laboratories. **11a** was dissolved in HS15/NaCl (4mg/mL, 7.5/100, v/v), and the **11a** was administrated once daily via intravenous drip. The SD rats were dosed 24 mg/kg (one rat), 40 mg/kg (ten rats) and 60 mg/kg (four rats) via intravenous drip administrated for acute toxicity study.

**Dose range toxicity study for one week of 11a**

SPF SD rats were half male and half female, which were weighted of 180~220 g for females and 200~240 g for males. Beagle dogs were half male and half female, which were weighted of 9-11 kg. Dose range toxicity studies for one week were performed in GLP laboratories. SD rats were assigned to four groups which contained one vehicle group and three intravenous drip administrated (0.2 mL/min) groups, the dosage of **11a** were 2, 6, 18 mg/kg (four mice per group), respectively. The Beagle dogs (four dogs) were dosed via intravenous drip administration (1.5 mL/min) (10 mg/kg (the first day), 15 mg/kg (the second day), 20 mg/kg (the third day), 25 mg/kg (the fourth day), 25 mg/kg (the fifth to seventh days, randomly two dogs), 40 mg/kg (the fifth to seventh days, other two dogs)). All animals were clinically observed once a day at least during 7 days for toxic signs which including bodyweight, food intake, and hematology. At the end of the experiment, samples of heart, liver, spleen, lung, kidney and administration site were collected.

All procedures relating to animal handling, care, and treatment were performed according to the guidelines approved by the Institutional Animal Care and Use Committee of the contract research organizations performing the study.

**Table S4. In *vivo* toxicity study of 11a**

| Study | Administration | Species | Dosage (mg/kg) | Frequency | Results |
|---|---|---|---|---|---|
| **Acute toxicity** | intravenous drip (0.2 mL/min) | SD rats | 24 mg/kg<br>40 mg/kg<br>60 mg/kg | Single-dose | No rats died after receiving 24, 40 mg/kg,<br>and one rat died after receiving 60 mg/kg |
| **Dose range studies for one** | intravenous drip (0.2 mL/min) | SD rats | 2 mg/kg<br>6 mg/kg<br>18 mg/kg | Repeat dose | No obvious toxicity<br>1. During the administration period, no anomalies of weight and general state were observed |

| | | | | | |
|---|---|---|---|---|---|
| **week** | | | | | in each group.<br>2. At the end of administration, the rats in each group underwent hematological and biochemical examination, and no anomalies were observed.<br>3. At the end of the administration, histological examination of the heart, kidney and lung were conducted, no anomalies were observed in each group |
| | intravenous drip (1.5 mL/min) | Beagle dogs | 10 mg/kg (the first day)<br>15 mg/kg (the second day)<br>20 mg/kg (the third day)<br>25 mg/kg (the fourth day)<br>25 mg/kg (the fifth to seventh days, randomly two dogs)<br>40 mg/kg (the fifth to seventh days, other two dogs) | Dose escalation | No obvious toxicity<br>1. When administered at a dosage of 25 mg/kg or more, the skin of the extremities (two of four dogs) developed allergic symptoms (lumpy, transient and recovery on the same day) during the administration period.<br>2. At the end of administration, there were no anomalies in hematology, blood biochemistry.<br>3. At the end of the administration, histological examination of the heart, lung, kidney, spleen and liver were conducted, no anomalies were observed in each group. |
